# An inhibitory feedback circuit mediating sleep need and sensory gating in *Drosophila*

**DOI:** 10.1101/2025.11.17.688806

**Authors:** Cedric B. Brodersen, Johannes Wibroe, Lorna May Shakespeare, David Owald, Katharina Krohn, Davide Raccuglia

**Affiliations:** Institute of Neurophysiology, Charité – Universitätsmedizin Berlin, corporate member of Freie Universität Berlin and Humboldt-Universität zu Berlin, Charitéplatz 1, 10117 Berlin, Germany

## Abstract

Many animals integrate sensory information during the day and sleep at night. How sensory processing during wakefulness contributes to making an organism tired at night, however, remains elusive. Here, we investigate the role of excitatory helicon cells in *Drosophila* (ExR1), a distinct neural population dedicated to processing sensory information. Using combined optogenetics and voltage imaging, we show that helicon cells excite sleep-promoting R3m ring neurons and trigger a subsequent autoinhibition of R3m via voltage- and Ca^2+^-gated K^+^ channels such as Slowpoke. Investigating the flies’ sleep patterns, we found that blocking synaptic output from helicon as well as RNAi-mediated knockdown of Slowpoke in R3m reduces sleep quality and leads to loss of nocturnal sleep, indicating that sleep need is generated via this route. Further, we show that excitation of R3m induces feedback inhibition of helicon cells via ionotropic GABA receptors. Knockdown of the ionotropic GABA receptor subunit Rdl in helicon cells increases sleep latency and causes sleep loss at the beginning of the night. We show that inhibition via Rdl facilitates nocturnal slow-wave activity which forms a sensory filter and prevents sleep disruption through auditory stimuli. We therefore uncover an inhibitory feedback circuit, in which neurons processing sensory information directly activate sleep promoting neurons to generate sleep need. At night, this sleep need is then converted into actual sleep, facilitated by the formation of a neural filter that stabilizes sleep/wake transition.

## Introduction

The basic concept of sleep homeostasis is that continuous neural activity during wakefulness generates sleep need that makes us feel tired (Borbely 1982). Sleep need, however, does not merely depend on the time spent in wakefulness, but is inextricably linked to sensory processing (Horne and Walmsley 1976; Engel-Yeger and Shochat 2012). This is particularly apparent in mental disorders such as schizophrenia and attention deficit/hyperactivity disorder, in which sensory overload easily exacerbates symptoms of irritability, restlessness and fatigue (Panagiotidi et al. 2020; Freeman and Waite 2025). In sleep deprived individuals, excessive sensory input can lead to synchronized electrical patterns within sensory neural networks which correlate with the occurrence of cognitive deficits (Quercia et al. 2018; Vyazovskiy et al. 2011). This shows that synchronized slow-wave activity for example is a neural marker for sleep need and creates sensory filters that regulate sensory processing to ensure adequate sleep quality (Krueger et al. 2008; Suárez-Grimalt and Raccuglia 2021). Precisely how neural networks processing sensory information are functionally linked to sleep-promoting neurons and how this interaction fine-tunes nocturnal sleep remains unclear.

To address this question, we focused on the *Drosophila* helicon cells (ExR1) which form a distinct population of four neurons with a high number of synaptic connections that radially project into large parts of the central complex (Hulse et al. 2021). The central complex is comprised of several different neural circuits that integrate multiple functions such as processing various sensory stimuli (Seelig and Jayaraman 2015; Fisher et al. 2019), locomotion control (Strauss and Heisenberg 1993), navigation and the representation of self-motion (S. S. Kim et al. 2017; Turner-Evans et al. 2017), providing the basis for action selection (Fiore et al. 2015; Sun et al. 2017). Within the central complex, the helicon cells receive homeostatic information from the dorsal fan shaped body, in which the accumulation of reactive oxygen species keeps track of the time spent in wakefulness (Jones et al. 2025; Kempf et al. 2019; Sarnataro et al. 2025). Together with R5 ring neurons helicon cells generate synchronized slow-wave activity that creates a nocturnal sensory filter to promote sleep quality, showing that these neurons directly operate at the intersection between sensory processing and sleep regulation (Raccuglia et al. 2025). Helicon cells are dedicated to processing visual information and gating locomotion, potentially also representing a multisensory integrational hub (Raccuglia et al. 2025). Prolonged optogenetic stimulation of helicon cells acutely increases locomotion (Donlea et al. 2018) followed by a subsequent sleep rebound, indicating that sensory information processed through the helicon cells could contribute to generating sleep need (Donlea et al. 2018). The mechanism of how synaptic output from this distinct neural population generates sleep need is unclear.

In this work, we uncover that synaptic output from helicon cells generates sleep need via an inhibitory feedback circuit with R3m ring neurons. Helicon cells activate R3m neurons and trigger an autoinhibition of R3m mediated by voltage- and Ca^2+^- activated K^+^ channels such as Slowpoke. We find that Slowpoke in R3m neurons contributes to generating sleep need. Moreover, we show that R3m ring neurons provide GABAergic feedback inhibition to helicon cells. In helicon cells, the GABA receptor subunit Rdl promotes nocturnal slow-wave activity to establish a sensory gate which stabilizes sleep/wake transitions and facilitates falling asleep at night. The inhibitory circuit represents a functional unit of how sensory processing during wakefulness contributes to promoting sleep quality at night.

## Results

### Synaptic output from locomotion gating helicon cells generates sleep need

Helicon cells are a neural population dedicated to processing visual information and gating locomotion (Donlea et al. 2018). To investigate the hypothesis that continuous processing of sensory information in these neurons generates sleep need, we chronically restrained synaptic output by expressing tetanus toxin (TNT) in helicon cells (**Fig. 1**). Tetanus toxin cleaves off synaptobrevin from synaptic vesicles and thus prevents fusion with the presynaptic membrane and subsequent release of neurotransmitters (Sweeney et al. 1995). In control flies, we expressed a version of the protein in which a point mutation renders it unfunctional (IMPTNT). Measuring sleep performance using *Drosophila* Activity Monitors (DAMs), we found that flies expressing TNT in helicon cells displayed a significantly altered sleep pattern (**Fig. 1A**) with increased daytime sleep and severely reduced nighttime sleep (**Fig. 1B**). As activation of helicon cells has shown to promote locomotion (Donlea et al. 2018; Raccuglia et al. 2025), impairing transmitter release from helicon should lead to decreased locomotor activity. However, the locomotor activity when flies were awake was not consistently altered (**Fig. S1A**), indicating that nocturnal sleep is not reduced because locomotion is affected, but because the fly’s sleep need is diminished.

**Figure 1.**
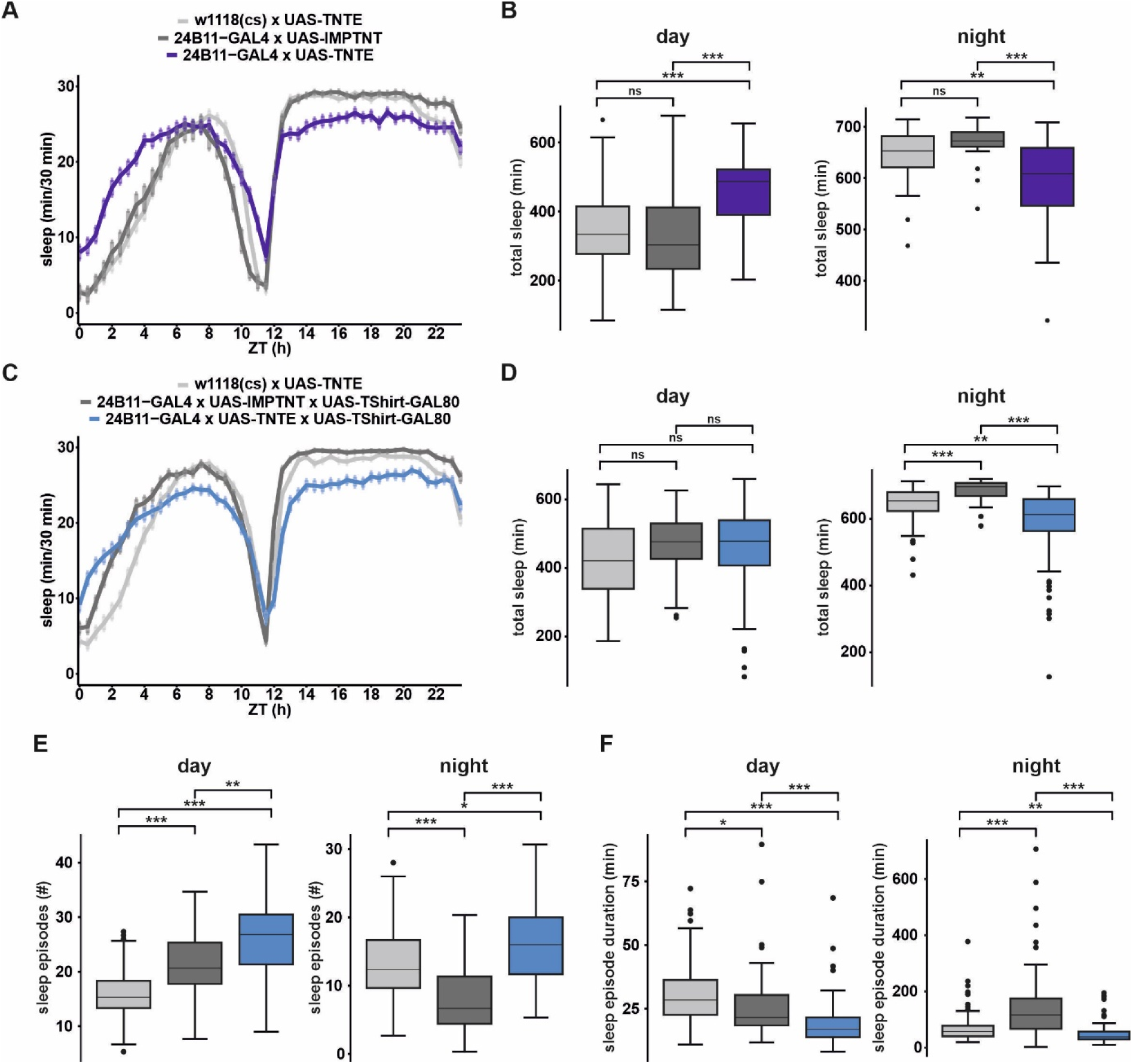
Blocking synaptic output from helicon cells reduces sleep quality and nocturnal sleep duration. **A** Average sleep pattern of flies expressing tetanus toxin (TNT) in helicon cells and respective controls expressing unfunctional TNT (IMPTNT) (n = 30-60) **B** Blocking helicon synaptic output (24B11-GAL4 x UAS-TNTE) increased total sleep during the day and decreased it at night (n = 30-60; Independent-Samples Kruskal-Wallis Test with Bonferroni correction). **C** Average sleep pattern of flies expressing TNT/IMPTNT and TShirt-GAL80, suppressing potential expression of TNT in the VNC (n = 66-81). **D** Blocking helicon synaptic output (24B11-GAL4 x UAS-TNTE x UAS-TShirt-GAL80) decreased total sleep duration at night (n = 66-81; Independent-Samples Kruskal-Wallis Test with Bonferroni correction). **E** Total number of sleep episodes is increased during day and at night (n = 66-81; Independent-Samples Kruskal-Wallis Test with Bonferroni correction). **F** Mean sleep episode duration is reduced during day and night (n = 66-81; Independent-Samples Kruskal-Wallis Test with Bonferroni correction).

To corroborate our findings, we additionally expressed a TShirt-GAL80 which inhibits expression of TNT within the ventral nerve cord (VNC) of *Drosophila* (Simpson 2016). Overall, the sleep pattern was similarly affected, showing strongly reduced nocturnal sleep (**Fig. 1C, D**). Although total sleep during the day was not significantly altered (**Fig. 1D**), both day and nighttime sleep were heavily fragmented with an increased number of sleep episodes (**Fig. 1E**) and a reduced sleep episode duration (**Fig. 1F**). As before, the fly’s locomotor activity when awake was unaffected by the genetic manipulation (**Fig. S1B**), substantiating the hypothesis that synaptic output from helicon cells creates a sleep need that is crucial to generate adequate sleep at night.

### Helicon cells induce sleep-regulating autoinhibition in R3m ring neurons

Helicon cells form reciprocal connections to two neural populations of ring neurons that have been shown to generate sleep need (R5) or directly promote sleep (R3m) (Liu et al. 2016; Raccuglia et al. 2019; Yan et al. 2023; Singh et al. 2023). As we have recently investigated the interaction between R5 and helicon and their role in regulating behavioral quiescence (Raccuglia et al. 2025), we here focused on the interaction between helicon and R3m ring neurons.

We used combined optogenetics and voltage imaging to investigate the functional connectivity between helicon cells and their sleep promoting synaptic partners, the R3m ring neurons (**Fig. 2**). First, we expressed the red-light activatable cation channel CsChrimson in helicon, while simultaneously expressing the genetically encoded voltage indicator ArcLight in R3m neurons (**Fig. 2A**). Immediately upon helicon cell activation, R3m neurons display a fast depolarization which is followed by a slower and longer lasting hyperpolarization (**Fig. 2B, C**). As helicon cells have been identified to be excitatory (Donlea et al. 2018) the initial depolarization of R3m is likely directly triggered by helicon. The hyperpolarization could either be a secondary network phenomenon or mediated cell-intrinsically by voltage- and Ca^2+^-activated K^+^ channels that mediate hyperpolarization upon excitation. To test for the latter, we applied the specific antagonist charybdotoxin (Miller et al. 1985) during optogenetic stimulation of helicon cells (**Fig. 2D**) and found that the hyperpolarization was significantly diminished while the depolarization of R3m was unaffected (**Fig. 2E**), indicating that indeed voltage- and Ca^2+^-activated K^+^ channels participate in mediating hyperpolarization observed in R3m.

**Figure 2.**
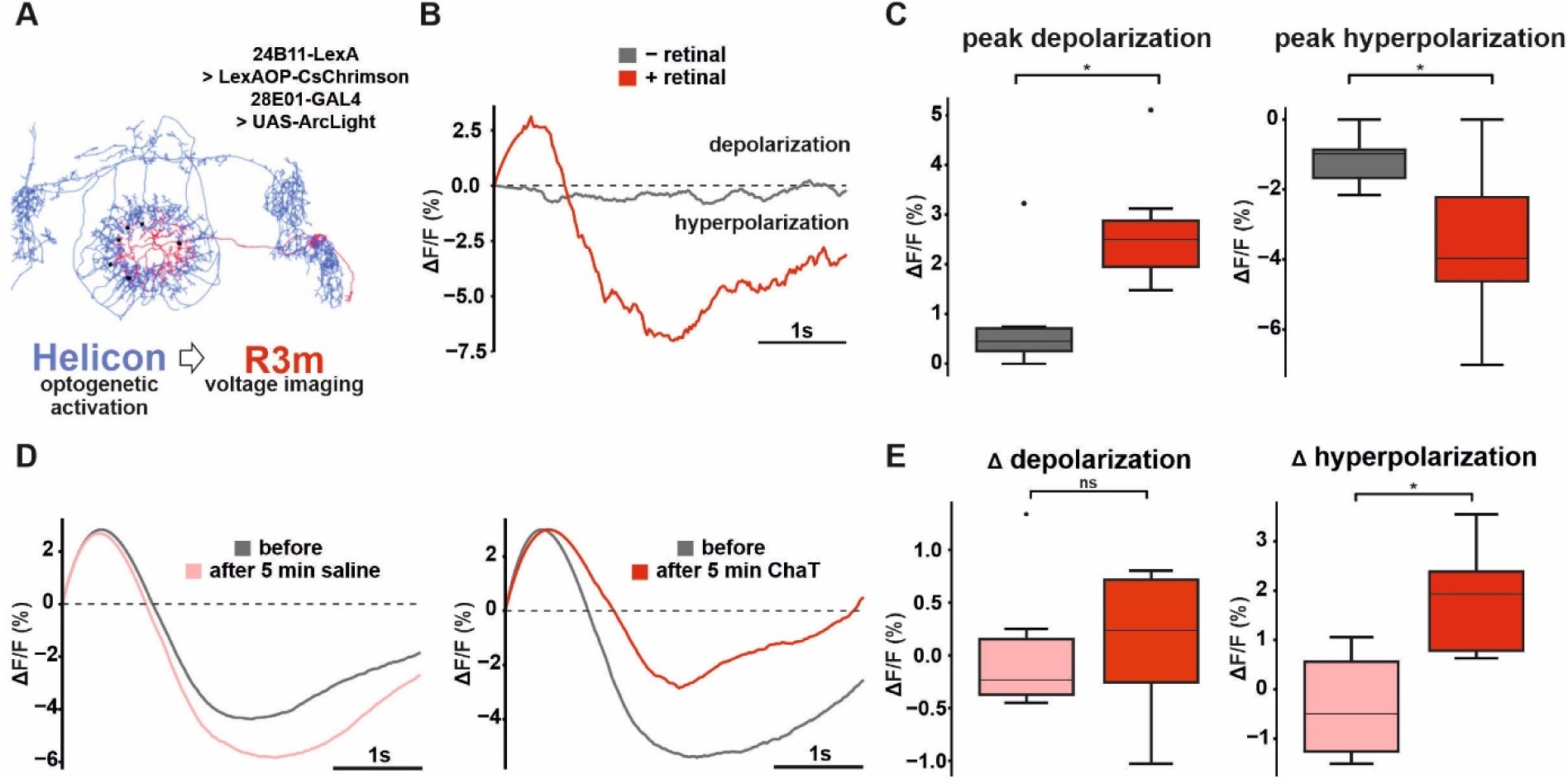
Helicon cells excite sleep promoting R3m neurons and trigger a hyperpolarization mediated by voltage- and Ca^2+^-gated K^+^ channels. **A** Connectomic reconstruction of a helicon cell (body-ID: 917647959) and a R3m ring neuron (body-ID: 1261768913). Black dots indicate synaptic connections from helicon to R3m. For concurrent optogenetics and voltage imaging, we expressed CsChrimson in helicon cells (24B11-LexA x LexAOP-CsChrimson) and ArcLight in R3m ring neurons (28E01-GAL4 x UAS-ArcLight). **B** Example electrical compound recordings of R3m neurons during optogenetic stimulation of helicon cells. Optogenetic stimulation requires the cofactor retinal. **C** Peak de- and hyperpolarization in R3m neurons during optogenetic stimulation of helicon cells (n = 6-8; Mann-Whitney-Test). **D** Example compound electrical recordings of R3m neurons during optogenetic stimulation of helicon cells before and after the application of the voltage- and Ca^2+^-gated K^+^ channel blocker charybdotoxin (ChaT) or saline (= control). **E** Charybdotoxin blocks helicon-triggered hyperpolarization in R3m neurons while the depolarization is unaffected (n = 6,8; Mann-Whitney-Test).

In *Drosophila*, Slowpoke is one of the most abundant voltage- and Ca^2+^-activated K^+^ channels (Atkinson et al. 1991). To test the impact of Slowpoke-mediated inhibition of R3m on sleep regulation, we performed an RNAi-mediated knockdown of *slowpoke*-expression using a specific split-GAL4 line to target R3m neurons (**Fig. 3**) and found that it led to loss of sleep specifically at night (**Fig. 3A, B**). Moreover, sleep quality was reduced due to partial sleep fragmentation, indicated by a reduced sleep episode duration (**Fig. 3C**) but an unaltered number of sleep episodes (**Fig. 3D**). Intriguingly, we found that locomotor activity when awake was reduced only during the night (**Fig. 3E**). This effect is not connected to the observed sleep fragmentation as a reduced locomotor drive would lead to increased and more consolidated sleep. This finding rather suggests that Slowpoke-mediated inhibition of R3m neurons contributes to gating locomotion and independently generates sleep need, potentially through interactions with their synaptic partners or through internal Slowpoke-mediated changes in gene expression.

**Figure 3.**
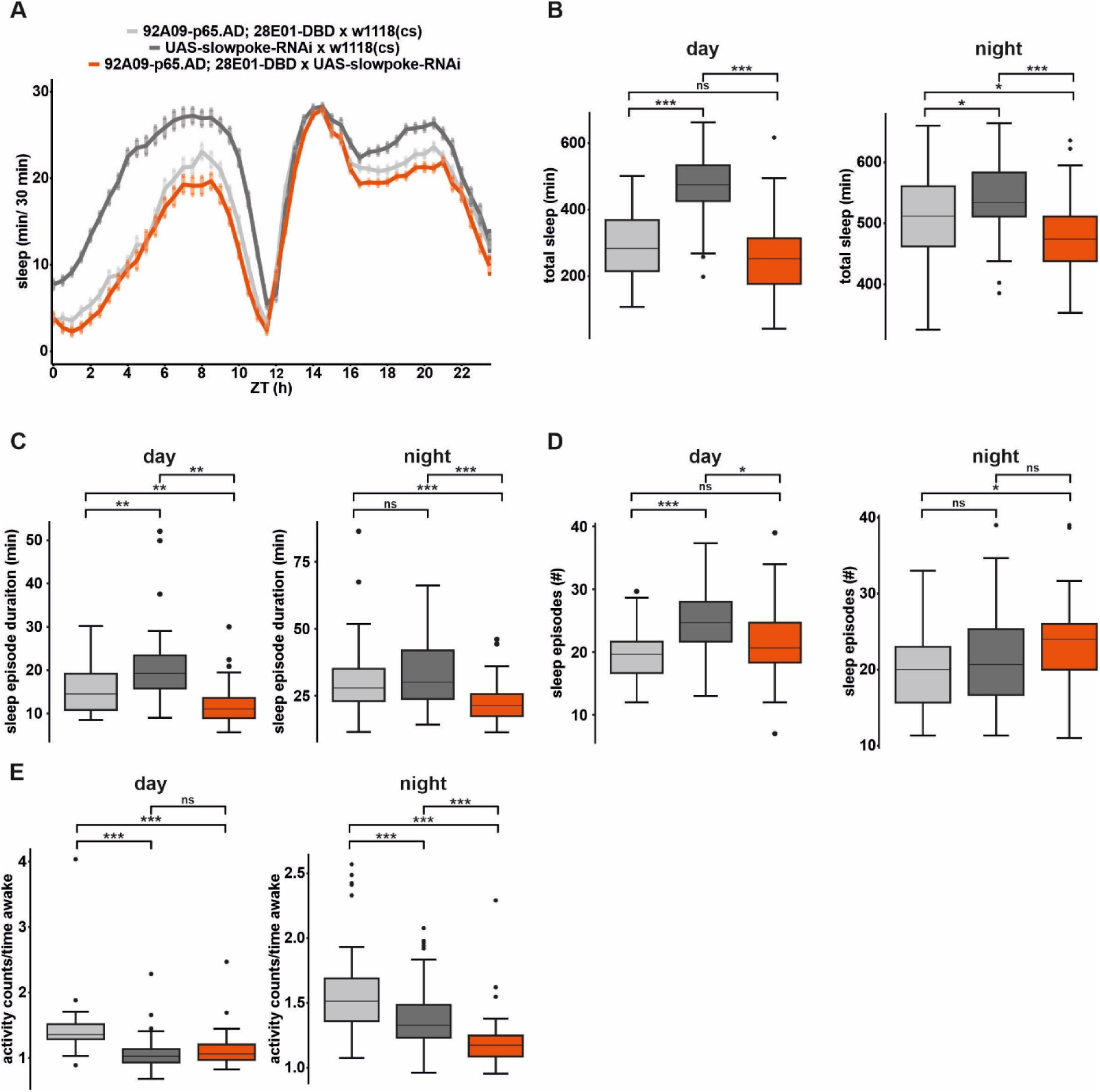
Knockdown of Slowpoke in R3m ring neurons leads to partial sleep fragmentation and nocturnal sleep loss. **A** Average sleep pattern of flies expressing Slowpoke-RNAi in a split-GAL4 driver line for R3m (92A09-p65.AD; 28E01-GAL4.DBD x UAS-Slowpoke-RNAi) and respective controls (n = 53-57). **B** Knockdown of Slowpoke reduces total sleep duration at night (n = 53-57, Independent-Samples Kruskal-Wallis Test with Bonferroni correction). **C** Mean sleep episode duration is diminished during day and at night (n = 53-57, Independent-Samples Kruskal-Wallis Test with Bonferroni correction). **D** Total number of sleep episodes is unaffected during day and at night (n = 53-57, Independent-Samples Kruskal-Wallis Test with Bonferroni correction). **E** Activity counts per time awake are reduced only at night. N = 53-57, Independent-Samples Kruskal-Wallis Test with Bonferroni correction).

Strikingly, our findings partially reproduce the sleep fragmentation and sleep loss observed when blocking synaptic output from helicon cells (**Fig. 1**), indicating that the generation of sleep need via helicon cells is linked to Slowpoke-mediated inhibition of R3m.

### GABAergic inhibition of helicon regulates sleep/wake transitions at the beginning of the night

Considering the reciprocal nature of their synaptic connectivity, we next aimed to investigate how R3m neurons feedback onto helicon (**Fig. 4**). We therefore inverted our previous experiment, now optogenetically stimulating R3m neurons while recording compound electrical activity in helicon (**Fig. 4A**). We found that activation of R3m neurons hyperpolarizes helicon cells (**Fig. 4B**). As R3m neurons are predominantly GABAergic (Singh et al. 2023), we tested whether ionotropic GABA receptors are involved in mediating this hyperpolarization. Indeed, we found that blocking ionotropic GABA receptors with picrotoxin reduced the R3m-mediated hyperpolarization of helicon (**Fig. 4 C, D**).

**Figure 4.**
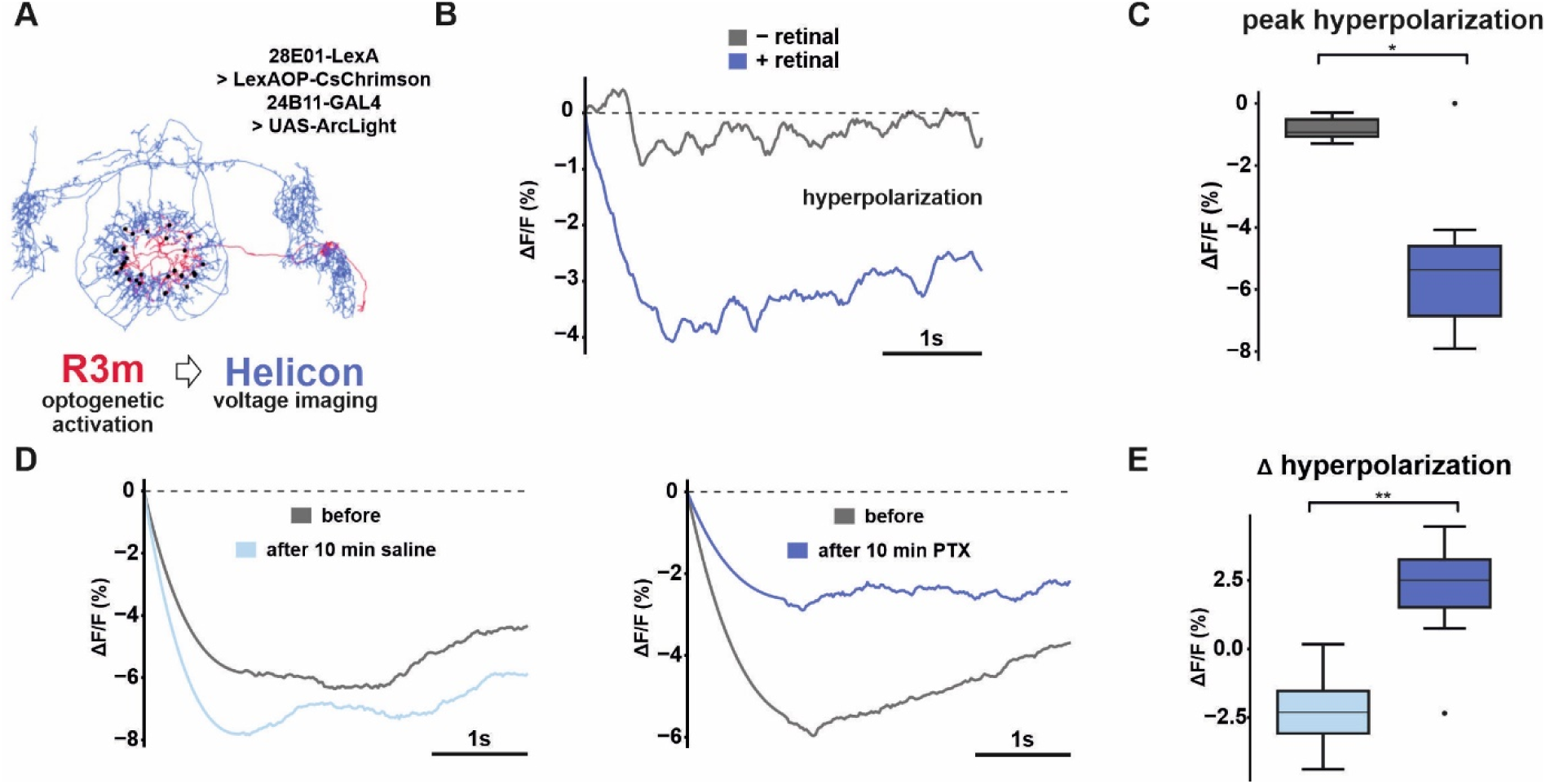
R3m ring neurons hyperpolarize helicon through GABAergic inhibition. **A** Connectomic reconstruction of a helicon cell (body-ID: 917647959) and a R3m ring neuron (body-ID: 1261768913). Black dots indicate synaptic connections from R3m to helicon. For concurrent optogenetics and voltage imaging, we expressed CsChrimson in R3m neurons (28E01-LexA x LexAOP-CsChrimson) and ArcLight in helicon cells (24B11-GAL4 x UAS-ArcLight). **B** Example electrical compound recordings of helicon cells during optogenetic stimulation of R3m. Optogenetic stimulation requires the cofactor retinal. **C** Peak hyperpolarization of helicon cells during optogenetic stimulation of R3m neurons (Mann-Whitney-Test; n = 5-7.) **D** Example compound electrical recordings of helicon cells during optogenetic stimulation of R3m before and after application of the Cl^-^ channel blocker picrotoxin (PTX, 100µM) or saline (as a control). **E** R3m-induced hyperpolarization of helicon cells is abolished by application of (Mann-Whitney-Test; n = 6-8).

In *Drosophila*, the most abundant subunit for ionotropic GABA receptors is Rdl (Lee et al. 2003). To investigate how GABAergic inhibition of helicon affects sleep need and quality, we employed RNAi-mediated knockdown of Rdl in helicon (**Fig. 5**) and found that sleep was reduced specifically during the first 4 hours of the night (**Fig. 5A, B**). Moreover, sleep latency was increased, meaning that flies needed longer to fall asleep at night (**Fig. 5C**). Locomotor activity when awake was not affected by the Rdl-knockdown (**Fig. S2A**). To corroborate our findings, we repeated the experiment with a specific split-GAL4 targeting the helicon cells (**Fig. 5D**). We were able to verify the observed behavioral phenotype and again found that sleep was reduced during the first 4 hours of the night (**Fig. 5E**) and nocturnal sleep latency was also increased (**Fig. 5F**). As in the previous experiment, locomotor activity when awake was not affected (**Fig. S2B**). Our findings indicate that ionotropic GABAergic inhibition of helicon regulates sleep/wake transitions at the beginning of the night.

**Figure 5.**
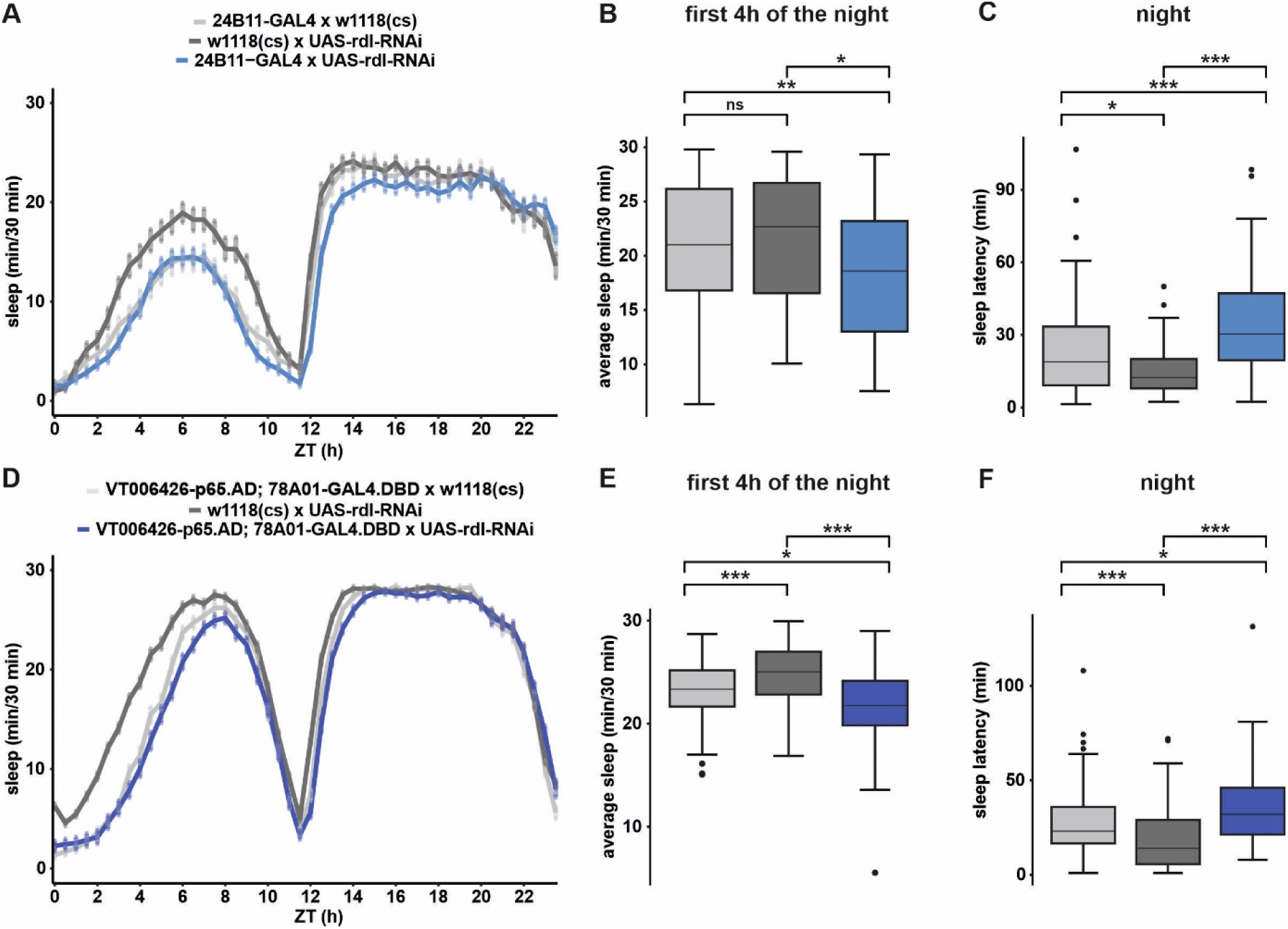
Knockdown of GABA receptor subunits Rdl in helicon cells diminishes the propensity to fall asleep at night. **A** Average sleep pattern of flies with knockdown of GABA receptor subunits rdl in helicon cells (24B11-GAL4 x UAS-Rdl-RNAi) and respective controls (n = 58-84). **B** Sleep duration during the first 4 hours of the night is reduced by knockdown of Rdl (n = 58-84, Independent-Samples Kruskal-Wallis Test with Bonferroni correction). **C** Sleep latency at night is increased by knockdown of Rdl (n = 58-84, Independent-Samples Kruskal-Wallis Test with Bonferroni correction). **D** Average sleep pattern of flies expressing Rdl-RNAi in a split-GAL4 line targeting helicon cells (V006426-p65.AD; 78A01-GAL4.DBD x UAS-Rdl-RNAi) and respective controls (n = 90-100). **E** Sleep duration during the first 4 hours of the night is reduced by knockdown of Rdl (n = 90-100, Independent-Samples Kruskal-Wallis Test with Bonferroni correction). **F** Sleep latency at night is increased by knockdown of Rdl (n = 90-100, Independent-Samples Kruskal-Wallis Test with Bonferroni correction).

### GABA receptor subunit Rdl facilitates helicon slow-wave activity and sensory filtering

Having trouble falling asleep might indicate that a filtering function within the brain is compromised. We have recently shown that helicon and R5 ring neurons generate synchronized slow-wave oscillations at night to establish a sensory filter that enables the fly to shut down the external world and facilitate behavioral quiescence (Raccuglia et al. 2025). To investigate whether slow-wave activity in helicon might be affected by Rdl-knockdown, we performed *in vivo* voltage imaging to measure the nocturnal compound electrical activity (**Fig. 6**). Indeed, we found that electrical oscillations between 0-1.5 Hz were significantly reduced (**Fig. 6A, B**) indicating that the sensory filter is not working optimally. To directly test this, we measured how easily sleep in male flies can be disrupted by presenting a male courtship song (**Fig. 6C-E**). For males, courtship song coming from other males signals a potential mating rival which can trigger aggressive behavior (Deutsch et al. 2019). We sleep-deprived flies over night and waited for them to fall asleep (5 min of quiescence) before presenting the auditory stimulus (**Fig. 6C**). Verifying increased sleep latency observed in the DAMs, knockdown of Rdl in helicon cells led to sleep-deprived flies needing longer to fall asleep in this experiment (**Fig. 6D**). When asleep, these flies woke up more readily than their controls, clearly showing that Rdl-knockdown in helicon compromises a sensory filter that shuts down the external world to promote transitions from wakefulness to sleep (**Fig. 6E**).

**Figure 6.**
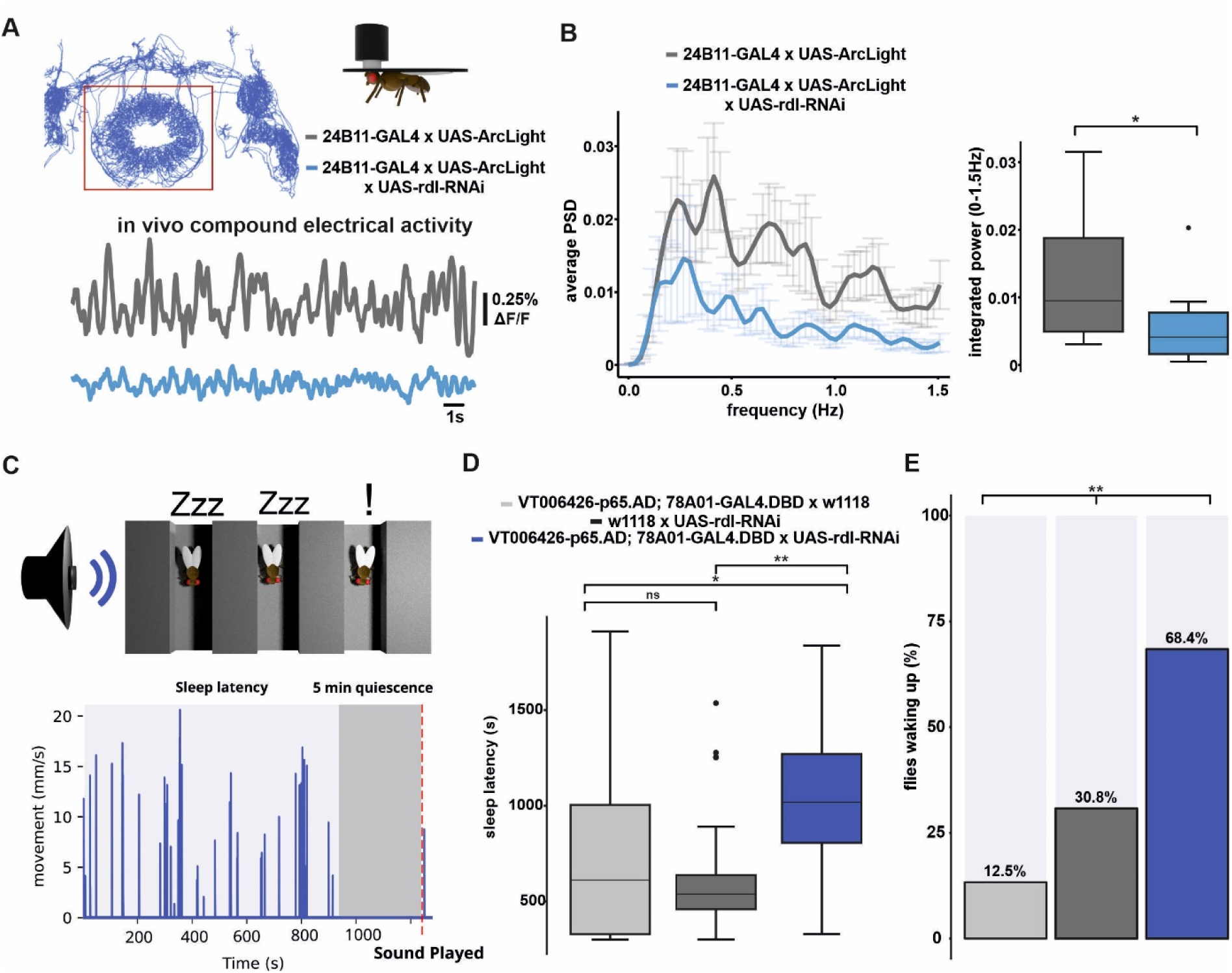
Knockdown of ionotropic GABA receptor subunit Rdl in helicon cells interferes with sensory filtering in tired and sleeping flies. **A** Connectomic reconstruction of all helicon cells indicating imaged synaptic connections. Example nocturnal compound electrical recordings (ZT 14-18) of helicon cells with knocked down GABA receptors Rdl (24B11-GAL4 x UAS-ArcLight x UAS-Rdl-RNAi) and respective control. **B** Average power spectrum (+/- SEM) and integrated power (0-1.5 Hz) showing a reduction of electrical slow-wave oscillations in helicon cells (Mann-Whitney-Test; n = 10-12). **C** Schematic showing experimental design in which sleeping male flies are presented with *Drosophila* courtship songs. Example activity profile of a fly showing the duration until the fly fell asleep (sleep latency) and 5 min of quiescence before the song was played. **D** Knockdown of Rdl in helicon cells (VT006426-p65.AD; 78A01-GAL4.DBD x UAS-Rdl-RNAi) increases sleep latency (n=15-26; Independent-Samples Kruskal-Wallis Test with Bonferroni correction). **E** Rdl-knockdown increases percentage of flies waking up to courtship song (n=15-26; Pearson Chi-Square Test).

## Discussion

To investigate how neural circuits dedicated to sensory processing generate sleep need, we used *Drosophila* and focused on a reciprocal connection between helicon cells and sleep promoting R3m ring neurons. We show that synaptic output from helicon cells is required for adequate sleep at night (**Fig. 1**). Further, we find that synaptic output from helicon cells generates sleep need independently of gating locomotion (**Fig. 1**). At least part of this sleep need is generated via activation of sleep-promoting R3m neurons which triggers a self-regulatory autoinhibition mediated by voltage- and Ca^2+^-gated K^+^ channels such as Slowpoke (**Fig. 2**). Knockdown of Slowpoke reproduces the nocturnal sleep loss observed after blocking synaptic output from helicon cells, suggesting that this neural route might compound sleep need during the day which leads to more sleep towards the end of the night (**Fig. 3**). The finding that sleep fragmentation from blocking helicon cell output is only partially reproduced by knockdown of Slowpoke in R3m indicates that helicon cells might also generate sleep need through targeting other synaptic partners, such as R5 ring neurons (Donlea et al. 2018). Previous work has shown that helicon cells excite R5 ring neurons which encode sleep need via NMDA receptor dependent plastic changes in electrical activity (Liu et al. 2016; Raccuglia et al. 2019). Therefore, R5 and R3m ring neurons might represent parallel routes that synergistically generate sleep need.

Our data implies that autoinhibition mediated via Slowpoke facilitates locomotion when flies are awake (**Fig.3**). This suggests that autoinhibition during wakefulness prevents feedback inhibition onto helicon cells and thus enables gating of locomotion (**Fig. 7**). Our data also indicates that synaptic output from the helicon cells generates sleep need by “charging” sleep promoting R3m neurons in a *slowpoke*-dependent manner. This could be mediated through network interactions but also cell-autonomously by Slowpoke-mediated changes in transcription and translation (E. Z. Kim et al. 2017). However, when sleep need is high at the beginning of the night, helicon cells start to display slow-wave activity and R3m neurons exert their sleep-regulating functions through GABAergic inhibition of helicon cells (**Fig. 5**), facilitating the physiological switch from the awake to the asleep state by hyperpolarizing helicon (Raccuglia et al. 2025) (**Fig. 7**).

**Figure 7.**
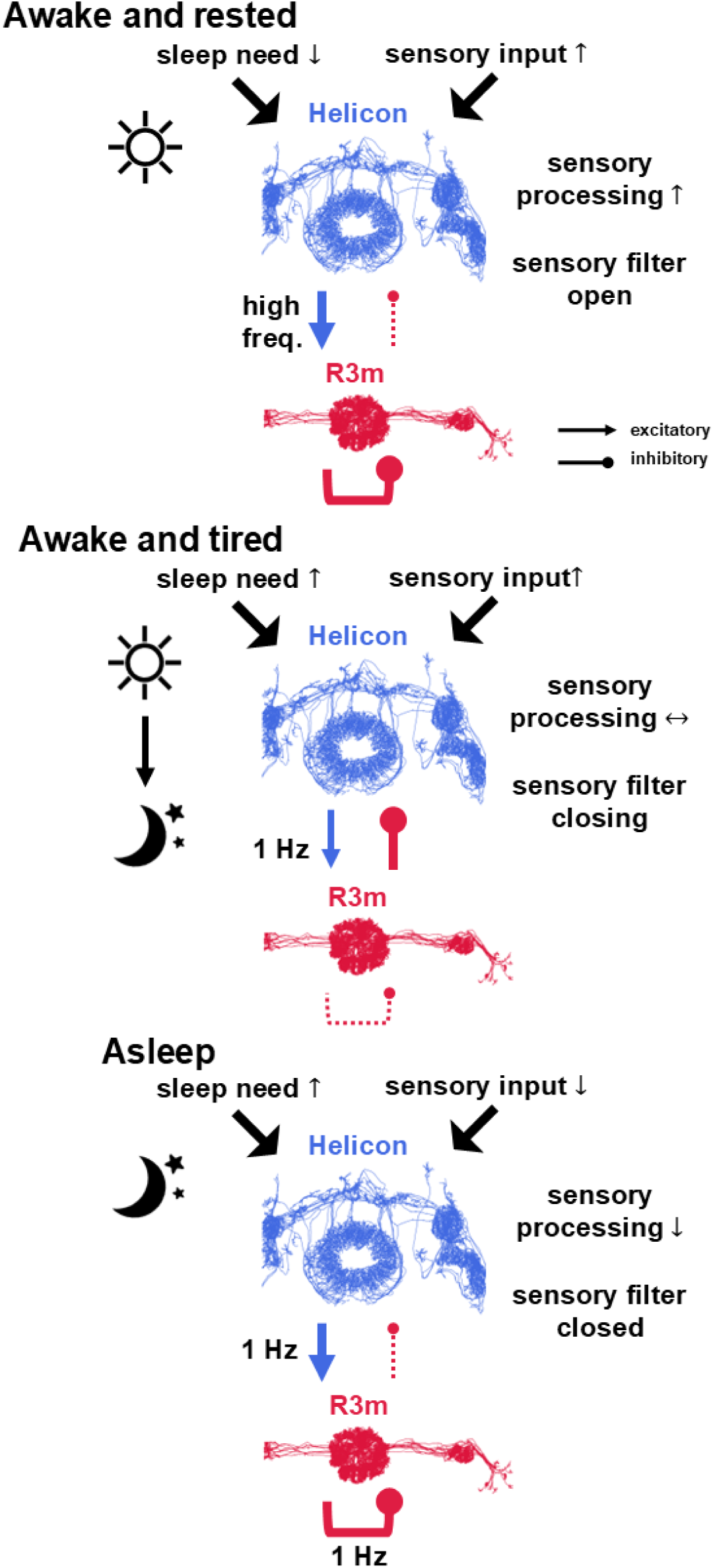
Graphical model illustrating how interactions between helicon and R3m might regulate state-dependent sensory gating. **Awake** During wakefulness helicon cells display high-frequency activity, exciting R3m neurons and triggering auto-inhibition of R3m which prevents feedback inhibition onto helicon cells. This keeps the sensory filter open and allows for sensory processing as well as gating locomotion through helicon cells. **Falling asleep** During the transition from day to night heightened sleep need leads to 1 Hz oscillatory activity which might facilitate a physiological switch from auto-inhibition of R3m towards increased feedback inhibition onto helicon cells. Hyperpolarization of the helicon cells further contributes to the formation of a sensory gate by increasing slow-wave activity and thus facilitates falling asleep by filtering sensory information. **Asleep** When asleep and helicon slow-wave activity is at its peak, it is transduced to R3m and causes a slow recurrent autoinhibition which effectively hyperpolarizes R3m. This stabilizes the sensory filter in support of maintaining behavioral quiescence.

Excitation of R3m neurons can trigger a feedback inhibition of helicon cells via ionotropic GABA receptors (**Fig. 4**). GABAergic inhibition through R3m is an important aspect in sleep regulation as the disruption of GABAergic neurotransmission in R3m neurons prevents post-injury sleep rebound (Singh et al. 2023). Here, we show that inhibition of helicon cells through ionotropic GABA receptors (Rdl) is particularly important during the transition between day and night, promoting sleep during the first hours of the night (**Fig. 5, 7**). We previously reported that lowering the membrane potential is required for helicon cells to generate slow-wave activity (Raccuglia et al. 2025). Driven by circadian input helicon cells synchronize their electrical patterns with R5 ring neurons to establish a sensory filter that facilitates nocturnal quiescence (Raccuglia et al. 2025). In this work, we show that GABAergic inhibition contributes to hyperpolarizing helicon cells and generating slow-wave activity that facilitates behavioral quiescence and thus promotes a physiological transition from wakefulness to sleep (**Fig. 5, 6**). Sleep in flies with knocked down Rdl receptors in helicon was more readily disrupted (**Fig. 6**), demonstrating a role for GABAergic input in establishing and maintaining a sensory filter for sleep (**Fig. 7**). Interestingly, the dFSB, which plays a role in homeostatic sleep regulation, also hyperpolarizes helicon cells, which has been shown to block visually- evoked activity (Donlea et al. 2018). Therefore, helicon cells integrate homeostatic and sensory information via two different inhibitory routes, both leading to an accumulation of sleep need and the creation of a neural filter that ultimately blocks sensory information to promote falling and staying asleep.

Our data indicates that Slowpoke is required to maintain nighttime sleep, particularly in the later phases of the night (**Fig. 3**). We propose that synchronous slow-wave activity between helicon and R5 might be transduced to R3m which triggers a recurrent auto-inhibition and prevents feedback inhibition onto helicon (**Fig. 7**). This would effectively stabilize synchronization and the sensory filter in support of maintaining behavioral quiescence.

In summary, we provide a neural framework that explains fundamental aspects of the reciprocity between sensory processing and sleep need. Neural networks convey sensory information to sleep promoting neurons and “charge” them across wakefulness, leading to the accumulation of sleep need. Together with circadian and homeostatic input, heightened sleep need promotes GABAergic inhibition of sensory neural networks. This inhibitory feedback motif establishes a selective sensory filter to inhibit sensory processing during sleep and provide sufficient sleep quality to alleviate sleep need. When sleep need is minimal at the start of a new day, the sensory filter opens again and allows for sensory information to be processed.

## Acknowledgements

We thank Jörg Geiger for providing the infrastructure that made this project possible. The authors also thank Eric Reynolds for comments on the manuscript and Anne von Philipsborn for providing the *Drosophila* courtship song. We also thank Lisa Scheunemann and Stephan Sigrist for providing several Drosophila stocks. Cedric B. Brodersen expresses his gratitude to Britta Eickholt and Christian Rosenmund for supporting his doctoral thesis. This project was funded by the Deutsche Forschungsgemeinschaft (DFG; German Research foundation; grant 462539941 to Davide Raccuglia).

---

**Figure S1.**
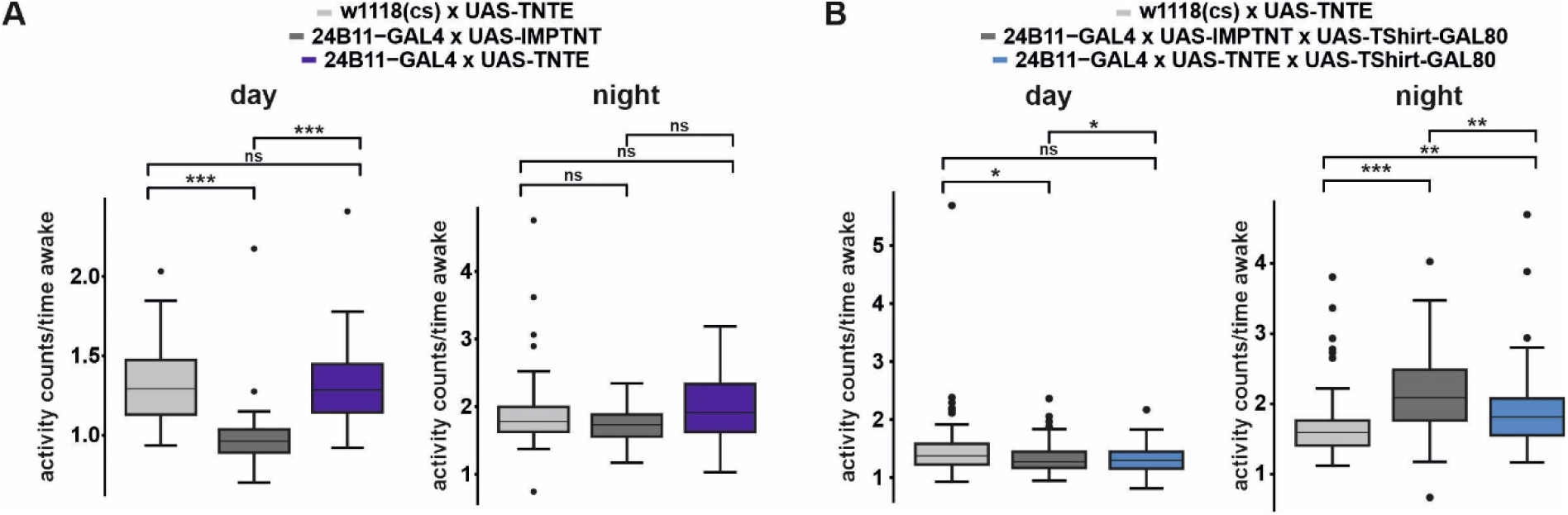
Blocking synaptic output from helicon cells leaves locomotor activity intact. **A** Blocking helicon synaptic output (24B11-GAL4 x UAS-TNTE) didn’t affect locomotor activity (n = 30-60; Independent-Samples Kruskal-Wallis Test with Bonferroni correction). **B** Blocking helicon synaptic output and additional expression of TShirt-GAL80 (24B11-GAL4 x UAS-TNTE x UAS-Tshirt-GAL80) did not affect locomotor activity. Please note that effects caused by genetic manipulation are considered relevant if the experimental group is consistently and significantly different from both genetic controls (see methods for details).

**Figure S2.**
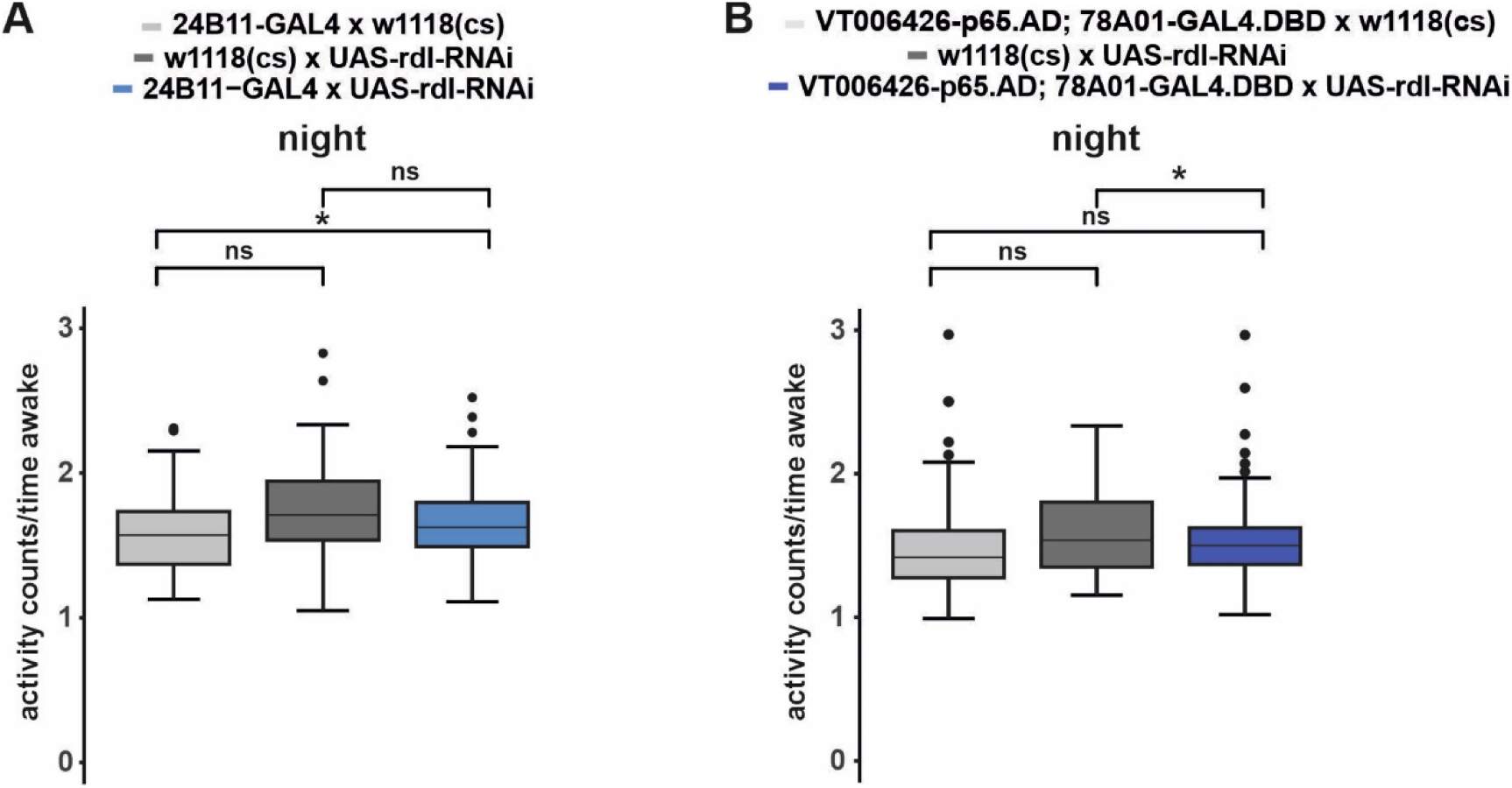
Rdl knockdown in helicon cells leaves locomotor activity intact. **A** Rdl knockdown in helicon cells (24B11-GAL4 x UAS-Rdl-RNAi) didn’t affect locomotor activity (n = 58-84; Independent-Samples Kruskal-Wallis Test with Bonferroni correction). **B** Rdl knockdown in a split-GAL4 line targeting helicon cells (V006426-p65.AD; 78A01-GAL4.DBD x UAS-Rdl-RNAi) didn’t affect locomotor activity (n = 90-100, Independent-Samples Kruskal-Wallis Test with Bonferroni correction). Please note that effects caused by genetic manipulation are considered relevant if the experimental group is consistently and significantly different from both genetic controls (see methods for details).

## Methods

### Flies

Flies were kept in plastic vials and reared on standard cornmeal food in incubators at 25°C at a humidity of 60% and under a 12h:12h light-dark cycle. Flies were obtained either from the Bloomington Stock Resource Center (BDRC), the Vienna Drosophila Resource Center (VDRC) or were gifts from specific labs. For details see Table 1.

**Table 1:**
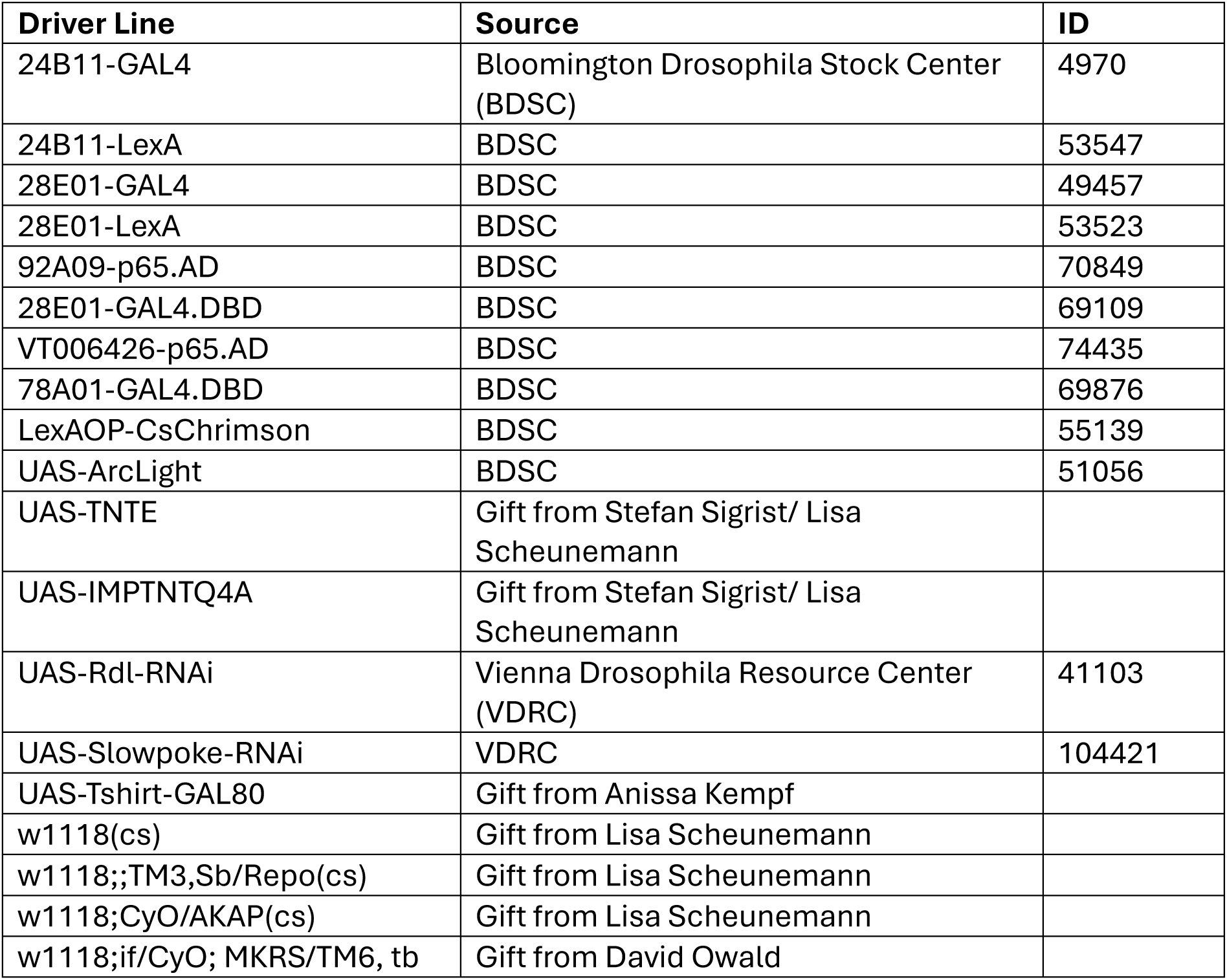
Sources of obtained *Drosophila* stocks.

### Drosophila activity monitors (DAMs)

Behavioral sleep analysis (**Fig. 1, 3, 5**) was performed using the DAM2 Drosophila activity monitors (trikinetics). Female non-virgin flies were collected 1-4 days after eclosion and then transferred into individual transparent tubes containing food on one side. After loading all tubes into the DAMs, experiments were performed in incubators at 25°C and 60% humidity. Experiments were conducted under a 12h:12h light-dark cycle in which lights were on between ZT 0 and ZT 12 (=day) and off between ZT 12 and ZT0 (night). Before starting the experiments, flies were allowed to acclimatize inside the tubes for 48 h. The flies’ sleeping pattern was recorded for 72 h and averaged to one 24-hour period using the DAMSystem3 software. In the average sleeping pattern, each datapoint corresponds to the minutes of sleep within a time period of 30 minutes (48 time periods over 24 hours). Using the DAMfilescan113X software recorded files were then converted into data showing the sleep profiles of individual flies. Activity traces of individual flies showing no movement were excluded from the analysis. We used the Vecsey Sleep and Circadian Analysis MATLAB Program v2 (SCAMP) for further analysis (Vecsey et al. 2024). The specific sleep and activity features (sleep latency, total sleep duration, mean sleep episode duration, number of sleep episodes and locomotion) were averaged separately for the day period, the night period and the full 24 hours. Analysis of the sleep amount within the first four hours (**Fig. 5**) of the night was performed by averaging the minutes of sleep per 30 minutes of the first eight 30-minute timeframes of the night.

To reduce potential genetic background effects, we backcrossed GAL4-lines (28E01-GAL4, 24B11-GAL4) as well as UAS-RNAi lines (*slowpoke* and *rdl*) 5 times with w1118(cs) wild type flies. Subsequently we crossed them with chromosome-specific balancer lines to create stable homozygous fly lines. Split-GAL4 AD and DBD lines were backcrossed 3 times during generating a stable fly line that carries both transgenes. For this, we used a double balancer line for chromosomes 2 and 3. UAS-TNTE and UAS-IMPTNT were not backcrossed as these 2 strains were already in the same genetic background and moreover, the expression of IMPTNT acts a specific genetic control for the expression of TNTE. To counteract residual genetic background effects, we only considered effects relevant in which the experimental group is consistently and significantly different from both genetic controls.

### Combined voltage imaging and optogenetics

For *ex-vivo* imaging experiments (**Fig. 2, 4**) we used 3-10 days old female flies and performed whole brain explant preparations as described previously (Cao et al. 2013). To block spontaneous activity ex vivo (**Fig. 2**), optogenetic experiments were conducted in a carbogenized high Mg^2+^ external solution consisting of (in mM) 70 NaCl, 3 KCl, 1.5 CaCl2, 20 MgCl2, 1 NaH2PO4, 10 glucose, 10 sucrose, 8 trehalose, 5 TES and 26 NaHCO3.

Depending on the strength of the GAL4 line we had to increase excitatory drive in some experiments (**Fig. 4**), by using a low (5 mM) Mg^2+^ external solution consisting of (in mM) 90 NaCl, 3 KCl, 1.5 CaCl2, 5 MgCl2, 1 NaH2PO4, 10 glucose, 10 sucrose, 8 trehalose, 5 TES and 26 NaHCO3. External solutions were adjusted to a pH of 7.4 with an osmolarity of approximately 280 mmol/kg. Charybdotoxin (Alomone labs) was used at a final concentration of 233 nM (**Fig. 2**) and Picrotoxin (Tocris) was used at 100 µM (**Fig. 4**).

For *ex-vivo* imaging we used an Olympus BX51W1 microscope and a Plan Apochromat 40x, numerical aperture 0.8, water immersion objective (Olympus). Recordings were performed at 50 frames per second using a Kinetix22 sCMOS camera (Teledyne Photometrics) controlled by micromanager.

For optogenetic experiments, we mixed all-trans-retinal (ATR) (Sigma) into the fly food as a 50 mM stock dissolved in 95% ethanol. The experimental flies were collected and transferred into vials with fly food containing 1.3 mM of all-trans-retinal 2-4 days before the experiment. For optogenetic stimulation CsChrimson was excited at a wavelength of 637/12nm while ArcLight was simultaneously excited at 473/35nm using a Lumencor Spectra X-Light engine (Lumencor) LED system. To avoid overexposure, we used an optical aperture to focally apply light, and LED power was adjusted for each experiment, making sure to use the minimum necessary light.

To assess peak de- and hyperpolarization (**Fig. 2, 4**) we determined the minimum and maximum changes in relative fluorescence within the first four seconds of the recording. To determine the changes in peak de- and hyperpolarization (**Fig. 2, 4**) we calculated the difference of the corresponding peaks in recordings performed before and after application of the drugs or saline (external solution) as controls.

### *In-vivo* imaging

In-vivo imaging (**Fig. 6**) was performed at ZT 14-18 (during a 12:12 h light:dark cycle the onset of light is at ZT 0 and the offset of light is at ZT 12) on 2-7 days old flies as previously described (Raccuglia et al. 2016). To mimic Mg^2+^ concentrations found in the *Drosophila* hemolymph, imaging was performed in the described 20 mM Mg^2+^ solution (Singleton and Woodruff 1994; Hofmann et al. 2010; Raccuglia et al. 2019). To prevent movements of the brain we applied 10% Papain (Roche) to the head capsule for 6 minutes and subsequently washed it out. Imaging was executed at a recording frequency of 27.785 Hz under an Olympus BX511W1 microscope with a 40x objective (NA 0.8, Olympus) using a IXon Ultra 897 camera (Andor) controlled by Solis I software (Andor). ArcLight was excited using a 470 nm LED controlled by LED controller DC4100 ( both from Thorlabs).

### Awakening sleeping flies with sound

For testing sleep depth and quality (**Fig. 6**), we used male flies between 3-10 days old that were sleep deprived overnight from 8 pm to 12 pm (16 h) in vials with standard food, 10-20 flies per vial, using a 2s mechanical stimulus randomised within a 20 second window. This was done with an analogue Multi-Tube Vortexer (VX-2500, VWR) controlled by TriKinetics acquisition software. Data was collected between ZT 3 and ZT 10. Behavioural data was collected using custom made closed-loop software written in python, which automatically presents a courtship song when sleep is detected (for details see https://github.com/johanneswibroe/multifruit). Sleep was defined as a period of quiescence of five minutes, and wakefulness was defined as a period with any detectable movement, including walking as well as grooming and wing motion. Auditory stimulus was a recording of a male pulse courtship song (gift from Anne von Philipsborn) modified to be 15 seconds long, played with a linear fade-in over 15 seconds, starting from silence and reaching a maximum level of 90 db SPL.

### Visualization, software, statistics

Each imaging recording was analyzed individually using the NOSA software applying background subtraction, baseline correction, smoothing algorithms and power spectrum analysis (Oltmanns et al. 2020). For visualization, each power spectrum was multiplied by 100. As the fluorescence of ArcLight increases with hyperpolarization and decreases with depolarization, traces were inverted to match common visualization. All data was statistically analyzed with IBM SPSS. In general, 2 groups were compared using the Mann-Whitney-Test, while more than 2 groups were compared using the Independent-Samples Kruskal-Wallis Test with Bonferroni correction. Binary values were compared using the Pearson Chi-Square Test. P-values are expressed as ns >0.05, * ≤0.05, **≤0.01, ***≤0.001. Data visualization was performed using RStudio with ggplot2 and CorelDraw2020 and 2024. Averaged sleep profiles and power spectrum are represented as mean +/- SEM. For boxplots we used the geom boxplot and stat boxplot functions in ggplot2 (Mcgill et al. 1978). (https://ggplot2.tidyverse.org/reference/geom_boxplot.html#references).

